# Fast calculation of small-angle scattering profiles of dense protein solutions modeled at the all-atom level

**DOI:** 10.1101/2025.07.14.664822

**Authors:** Sanbo Qin, Huan-Xiang Zhou

## Abstract

The small-angle scattering profile of a dense protein solution contains rich information about interprotein interactions, but extracting this information has been extremely challenging. In previous studies, we developed a fast Fourier transform (FFT)-based method, FMAPB2, to compute protein pair interactions at the all-atom level. Here, we use FMAPB2 to precompute pair interaction energies and then use the latter to drive simulations of dense protein solutions. The resulting MAEPPI (many-protein atomistic energy from pre-computed pair interaction) enables simulations of atomistic proteins at the speed of Lennard-Jones particles. On snapshots sampled from the simulations, we directly calculate the small-angle scattering profile. The results reveal artifacts generated by widely used spherical models and support a significant dimer population of bovine serum albumin at high concentrations. Our approach promises to modernize the prediction and interpretation of small-angle scattering profiles of dense protein solutions.

## Introduction

Small-angle scattering (SAS) is a uniquely powerful technique for probing the internal structures of proteins and interprotein interactions. Significant resources have been dedicated worldwide to collecting SAS data. These include X-ray beamlines at the Argonne National Laboratory ^1^, the European Molecular Biology Laboratory ^2^, Shanghai Synchrotron Radiation Facility ^3^, and NanoTerasu (Japan) ^4^, and neutron sources at the Oak Ridge National Laboratory ^5^, European Spallation Source ^6^, and Japan Proton Accelerator Research Complex ^7^. Most SAS studies were conducted at low protein concentrations (< 10 mg/mL) ^8^, where the data can be used to derive structural models or overall size measures such as the radius of gyration. At medium (10-40 mg/mL) and high (> 50 mg/mL) protein concentrations, SAS data contain information about weak interprotein interactions, which are the drivers in key biophysical problems including macromolecular crowding ^9^ and liquid-liquid phase separation ^10^. However, extracting this information from SAS data has remained a major challenge.

SAS experiments measure the dependence of the scattering intensity, *I*, on the magnitude,

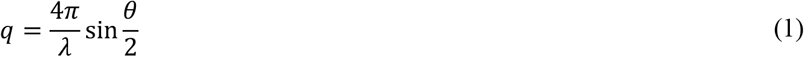

of the scattering vector, where *λ* is the wavelength of the incident beam and *θ* is the scattering angle. For a solution of a single protein species, if the protein molecules are approximated as being spherically symmetric, one can decompose the scattering intensity per unit sample volume into ^11,12^

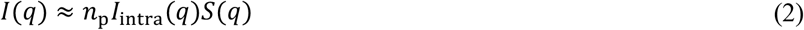

where *n*_p_ is the number density of the protein, *I*_intra_(*q*) is the scattering intensity of a single protein molecule [with normalization chosen such that *I*_intra_(0^+^) = 1; *q* = 0^+^ means taking the *q* → 0 limit], and *S*(*q*) is the interprotein structure factor. The latter is given by the Fourier transform of the center-center pair distribution function *g*(*R*),

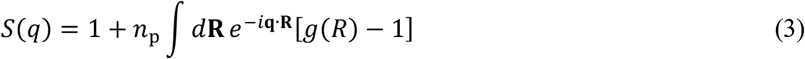

with **R** denoting the center-center displacement vector of a protein pair. Interprotein interactions control *g*(*R*) and hence the scattering profile, but extracting interprotein interactions from the scattering profile is daunting. Note that Eq (2) is exact only at *q* = 0^+^. In an ideal solution where intermolecular interactions are absent, *g*(*R*), *S*(*q*), and *I*(0^+^)/*n*_p_ all equal 1. For arbitrary *q*, it is sometimes helpful to define an effective structure factor, *S*_eff_(*q*), in analogy to Eq (2):

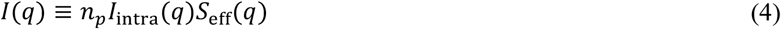

To obtain *S*_eff_(*q*) from *I*(*q*), a common practice is to use *I*_intra_(*q*) calculated by modeling the protein molecule as an ellipsoid with a uniform scattering length density ^13-17^, although *I*_intra_(*q*) can be determined from scattering experiments under dilute conditions or calculated from a crystal structure of the protein ^18-27^ [also http://debyer.readthedocs.org/; https://github.com/Niels-Bohr-Institute-XNS-StructBiophys/CaPP/].

Up to now, almost all quantitative analyses of SAS data under non-dilute conditions are based on Eq (2), i.e., the spherical approximation (or a slight variation thereof). This approximation is poor for anisotropic proteins such as antibodies ^28^. In any event, to calculate the structure factor *S*(*q*), one often models interprotein interactions by a spherically symmetric potential and invokes a further approximation, in the form of a closure relation ^13-15,17,29-31^. Closure approximations may incur significant errors, such that the resulting *S*(*q*) when used to fit experimental *S*_eff_(*q*) leads to unreliable estimates for the parameters in the interaction potential ^32^. Based on Eq (3) (or an equivalent expression), molecular simulations have been used to calculate *S*(*q*) for spherical ^32^ as well as more complicated models of interprotein interactions, including both coarse-grained ^28,33^ and all-atom models ^16,34^. A modified random-phase approximation (mRPA) was recently introduced to obtain *g*(*R*) and *S*(*q*) for all-atom models ^12^.

We now present an approach for performing fast simulations of dense protein solutions at the all-atom level and directly calculating *I*(*q*) from these simulations. Fast simulations are made possible by precomputing interaction energies of protein pairs. Precomputing of pair interactions has recently used to drive coarse-grained simulations ^35^. Our all-atom interaction energies of protein pairs are obtained using a method called FMAP (fast Fourier transform-based modeling of atomistic protein-protein interactions), where interaction energies are expressed as correlation functions and evaluated via fast Fourier transform (FFT) ^36,37^. A version of FMAP dubbed FMAPB2 has been used to determine second virial coefficients ^38^ and implement mRPA ^12^. Here the resulting MAEPPI (many-protein atomistic energy from pre-computed pair interaction) enables simulations of atomistic proteins at the speed of Lennard-Jones particles. *I*(*q*) directly calculated from MAEPPI simulations reveals additional shortcomings of spherical models and leads to a better understanding on the effects of interprotein interactions, protein concentrations, and preformed dimers and higher oligomers.

## Results

### MAEPPI: <u>m</u>any-protein <u>a</u>tomistic <u>e</u>nergy from <u>p</u>re-computed <u>p</u>air <u>i</u>nteraction

We explicitly represent the *n* atoms of each protein molecule. Assuming that the protein is rigid, the atomic positions are fully specified by the center position and orientation of the protein molecule. Our model for the interaction energy of a protein pair is a sum of terms over all atom-atom pairs between the two molecules. Each atom-atom pair has three terms, representing steric repulsion, nonpolar attraction, and electrostatic interaction ^37,38^. The interaction energy of two protein molecules labeled 1 and 2 is denoted as *U*_12_, which is fully determined by the center-center displacement vector **R** and the relative orientation **Ω**. We specify **R** by its Cartesian components (*X, Y, Z*) in a body-fixed frame of molecule 1, and **Ω** by a rotation matrix 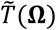, expressed in terms of three Euler angles, that transforms the body-fixed frame of molecule 1 to the body-fixed frame of molecule 2. For a system of *N* protein molecules, the total interaction energy is a pairwise sum:

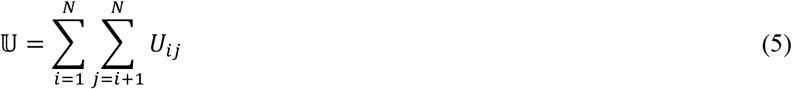

We precompute *U*_12_ for various **R** and **Ω**. For a given **Ω**, a single FMAPB2 calculation generates the values of *U*_12_ at all points on a cubic grid (with a 0.6-Å spacing; Figure 1A, each row). We then repeat the calculations for a set of **Ω** values that uniformly sample the rotational space ^39^ (Figure 1A). (We tested set sizes of 72, 576, and 4608, corresponding to 60º, 30º, and 15º separations between successive rotations.) These precomputed *U*_12_ values are then stored as a table (Figure 1B). With 310 grids per dimension of **R** and 72 rotations for **Ω**, the size of the table is only 2 GB in RAM.

**Figure 1.**
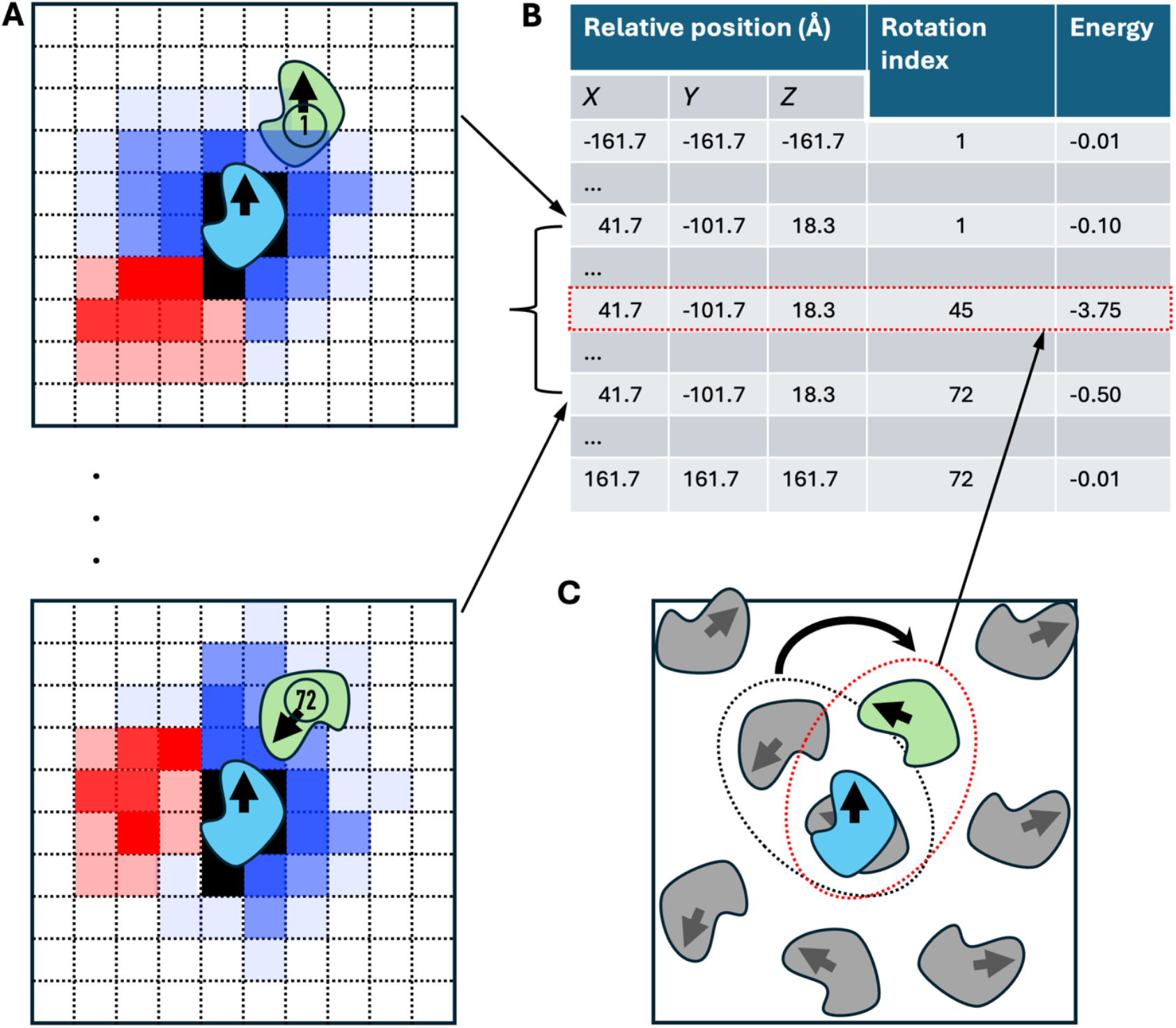
Precompution of protein pair interactions and their retrieval during MAEPPIsim. (A) For each relative orientation between a pair of protein molecules, the interaction energies at relative positions on a cubic grid are calculated using FMAPB2. (B) The interaction energies for all relative positions and orientations are saved to a table. (C) During MAEPPIsim, the interaction energy for each protein pair is obtained by first finding their relative position and orientation and then retrieving the energy in the corresponding entry from RAM.

In a many-protein system, the position and orientation of molecule *i* are specified by the center position **R**_)_ and the rotation matrix 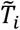 ; the latter yields the present orientation of the molecule when applied to its orientation in its body-fixed frame. Then the displacement vector between molecules *i* and *j* is

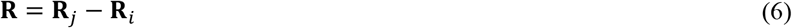

and the rotation matrix specifying their relative orientation is (Figure 1C)

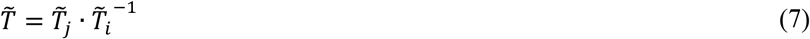

The interaction energy *U*_*ij*_ can then be found from the entry corresponding to **R** and 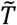 in the prestored table (Figure 1B, C). Enumeration over all protein pairs then yields the total interaction energy, which we call MAEPPI.

### MAEPPIsim: MAEPPI-enabled simulations

We use MAEPPI to drive Monte Carlo simulations, with moves in Cartesian coordinates and Euler angles. The default number of distinct rotations in the precomputed table is 72. MAEPPIsim takes only ∼10% longer than corresponding simulations of Lennard-Jones particles (Figure S1; see Table S1 for parameter values). For an 800-protein system in a cubic box with a 984-Å side length, representing lysozyme at 20 mg/mL, MAEPPIsim completes 20,000 Monte Carlo cycles in 14 min on an Intel Xeon Platinum 8358 2.6 GHz 32-core processor (Figure 2A). With a higher number of rotations covering the rotational space, the size of the precomputed table in RAM increases in proportion, but the simulation speed is virtually unaffected. We calculated various properties using 72, 576, and 4608 rotations; they all point to the conclusion that 72 rotations are sufficient for accuracy.

**Figure 2.**
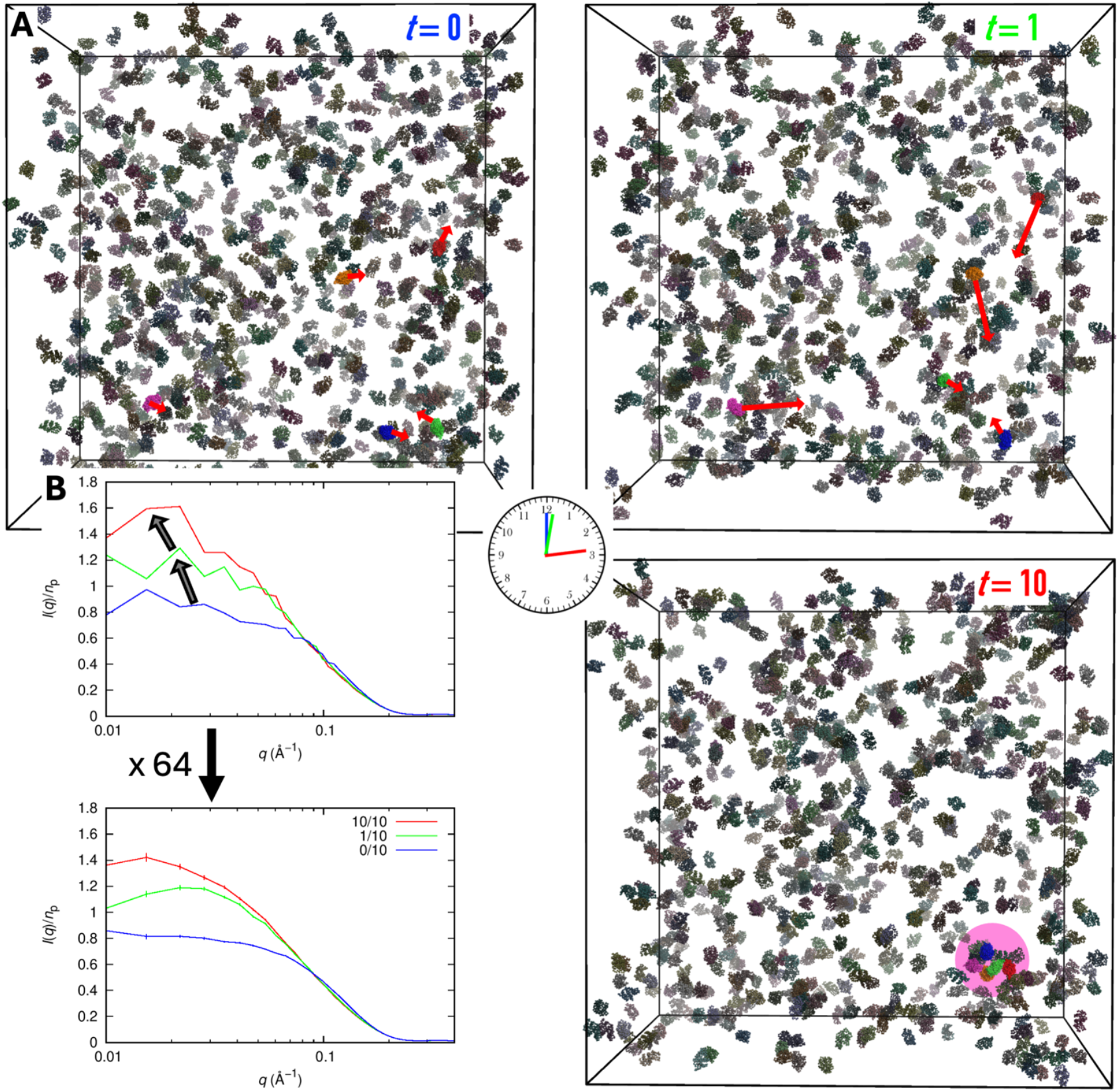
An example of MAEPPIsim + Sim2Iq. (A) Three snapshots from a simulation, at 0, 2,000, and 20, 000 Monte cycles (labeled as *t* = 0, 1, and 10). Five protein molecules that eventually form a cluster are in color; their movements are indicated by red arrows. Other molecules are in gray. The simulation box has a side length of 984 Å and contains 800 lysozyme molecules (corresponding to 20 mg/mL). Solvent conditions: 10 ºC and 0.15 M ionic strength. (B) Top: Sim2Iq result from the three snapshots in (A); bottom: average from 64 replicate simulations, with standard error of the mean shown as error bars.

As a simple test, in Figure S2A we compare the center-center pair distribution functions for 20 mg/mL lysozyme calculated from simulations using 72, 576, and 4608 rotations; the results are indistinguishable. Using the final snapshots of such simulations, we performed Widom insertion and calculated the interaction energy of a test protein with the protein solution in two ways. The first was by the FMAP method, which gives the interaction energies when the test protein was placed at all points on a cubic grid ^36,37^; the second was using the MAEPPI approach, where the interaction energy was read from the precomputed table as if in MAEPPIsim. In Figure S2B, C, we compare the FMAP and MAEPPI energies at the grid points by presenting the histograms of the energy pairs. The histograms have a very sharp peak on the diagonal (i.e., equal values from FMAP and MAEPPI), and a symmetric shape with respect to the diagonal. The symmetric shape produces error cancellations when performing averages over configurations. The spread in the off-diagonal direction is only moderately greater than that when comparing FMAP energies with the exact values from atom-based enumeration ^37^. When benchmarked against FMAP, MAEPPI results calculated using 72 and 4608 rotations are very close to each other, with only slight underpopulation of the peak region (Figure S2B inset, black arrowhead) and a slightly higher number of off-diagonal outliers (Figure S2B, orange arrowhead).

The sum of the Boltzmann factors of the interaction energies yields the excess chemical potential (*μ*^ex^) ^40^. Both the full *μ*^ex^ and its steric component show good convergence among simulations with 72, 576, and 4608 rotations (Figure S2D). These values also agree with those calculated by FMAP.

### Sim2Iq: from simulations to *I*(***q***)

The scattering intensity of a protein solution, after buffer subtraction, is given by

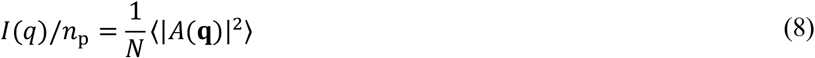

where ⟨⋯ ⟩ signifies an average over protein configurations. The scattering amplitude is

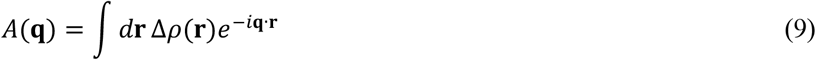

where Δ*ρ*(**r**) is the excess electron density (i.e., difference in electron density between the protein solution and buffer). Following Grant ^26,27^, we write Δ*ρ*(**r**) as

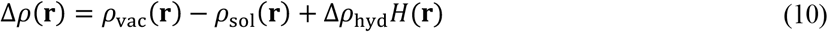

where the three components represent the electron density of the protein molecules when placed in vacuum, the electron density of the would-be solvent in the region excluded by the protein molecules, and the difference in electron density between hydration and bulk water in the hydration layer [represented by the function *H*(**r**)]. We map Δ*ρ*(**r**) to a cubic grid (Figure S3) and then perform an FFT to obtain *A*(**q**), and realize ⟨⋯ ⟩ by averaging over the last snapshots from 64 replicate simulations (Figure 2B).

For *ρ*_vac_(**r**) (Figure S3C), each atom makes an additive contribution, which takes the Cromer-Mann form ^41^:

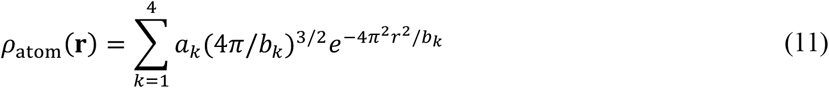

The sum of *a*_*k*_ is equal to the number of electrons in the atom; to the original *b*_*k*_, we also added a Debye-Waller factor to account for positional fluctuations of the atom. We also model *ρ*_sol_(**r**) as a sum of atomic contributions (Figure S3D); the latter assumes a Gaussian form

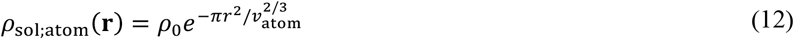

which, when integrated over **r**, yields a total of *ρ*_0_*ν*_atom_ electrons, with *ρ*_0_ denoting the electron density of bulk water and *ν*_atom_ denoting the solvent volume excluded by the atom. Lastly, the hydration-layer function *H*(**r**) is defined as the region between the bare van der Waals volume and an inflated van der Waals volume; for the latter, atomic radii are increased by 2.8 Å (Figure S3E). We refer to the *I*(*q*) calculation on snapshots sampled from MAEPPIsim as Sim2Iq. The default spacing for mapping Δ*ρ*(**r**) to a cubic grid is 1 Å. For an 800-molecule lysozyme system (a total of 1,568,00 atoms) in a cubic box with a 984-Å side length, one Sim2Iq calculation takes 12 s on an Intel Xeon Platinum 8358 2.6 GHz 32-core processor. The results for *I*(*q*) and *S*_eff_(*q*) are virtually unchanged when the spacing is reduced to 0.5 Å (Figure S4A, B). Also, neglecting the hydration-layer component of Eq (10) increases *I*(*q*) at intermediate *q* (around 0.1 Å^-1^; Figure S4C) but has little effect on *S*_eff_(*q*) for a repulsive condition (0.016 M ionic strength and 25 ºC) and medium concentration (20 mg/mL; Figure S4D). These effects can be attributed to a change in *I*_intra_(*q*) due to a decrease in the effective protein size by the neglect of the hydration layer. Further neglect of the second component in Eq (10) (compensated by an appropriate normalization) leads to a small decrease in *I*(*q*) at *q* > 0.3 Å^-1^.

As one more test of whether MAEPPIsim with 72 rotations is sufficient for accuracy, we compare in Figure S5A *I*(*q*) results calculated from simulations using 72, 576, and 4608 rotations. The close agreement attests the accuracy of using 72 rotations. We also tested the effect of varying the number of protein molecules in representing a protein solution at a given concentration. For 20 mg/mL lysozyme, the *I*(*q*) results calculated from simulations using *N* = 200, 400, and 800 molecules are compared in Figure S5B-D. The three sets of results agree well; however, *N* does affect the minimum *q* value (*q*_min_) at which *I*(*q*) can be calculated. *q*_min_ scales with the side length, *L*_box_, of the simulation box as *q*_min_ = 2.83π/*L*_box_. At a given concentration, a smaller *N* requires a smaller *L*_box_ and hence leads to a larger *q*_min_. The typical experimental *q*_min_ of 0.01 Å^-1^ corresponds to *L*_box_ = 890 Å.

Figure 2A presents the snapshots of a simulation at the start (*t* = 0), after 2,000 Monte Carlo cycles (*t* = 1) and 20,000 Monte Carlo cycles *t* = 10). The snapshot at *t* = 0 was prepared by random placement, and hence there was little chance for clustering of the protein molecules. At *t* = 1, the molecules started to form “dynamic” dimers and trimers due to mild attraction at the chosen solvent condition; at *t* = 10, even larger clusters formed. By “dynamic”, we mean that oligomers rapidly assemble and disassemble. Correspondingly, *I*(*q*)/*n*_p_ values at low *q* gradually moved up as larger clusters formed (Figure 2B). The *I*(*q*) curve from a single simulation was relatively noisy, but after averaging over 64 replicates, the curve became smooth, with relatively small error bars.

### *I* (*q*) results for lysozyme

In Figures 3A and S6, we compare the Sim2Iq result for *I*_intra_(*q*) of lysozyme (using Protein Data Bank 1AKI structure as input) and the experimental counterpart, deposited in SASBDB entry SASDU4 ^42^, showing excellent agreement. On the other hand, *I*_intra_(*q*) calculated using another code, Debyer (http://debyer.readthedocs.org/), shows overestimation around *q* = 0.1 Å^-1^. The same overestimation carries over to *I*(*q*)/*n*_p_ calculated at 20 mg/mL (Figure S7A), and is reminiscent of the *I*(*q*)/*n*_p_ result from Sim2Iq when only the vacuum component of the excess electron density is included (Figure S4C). Indeed, Debyer accounts only for the vacuum component, and agrees well with the vacuum-only Sim2Iq (Figure S7B). Debyer evaluates the scattering amplitude *A*(**q**) in direct space whereas Sim2Iq does so in Fourier space; the matching in results between the two methods thus provides strong cross-validation. We note in passing that the ellipsoid model severely underestimates *I*_intra_(*q*) at *q* > 0.3 Å^-1^(Figure S6), as already noted ^14^, and thus, when used to deduce *S*_eff_(*q*), can lead to significant errors.

**Figure 3.**
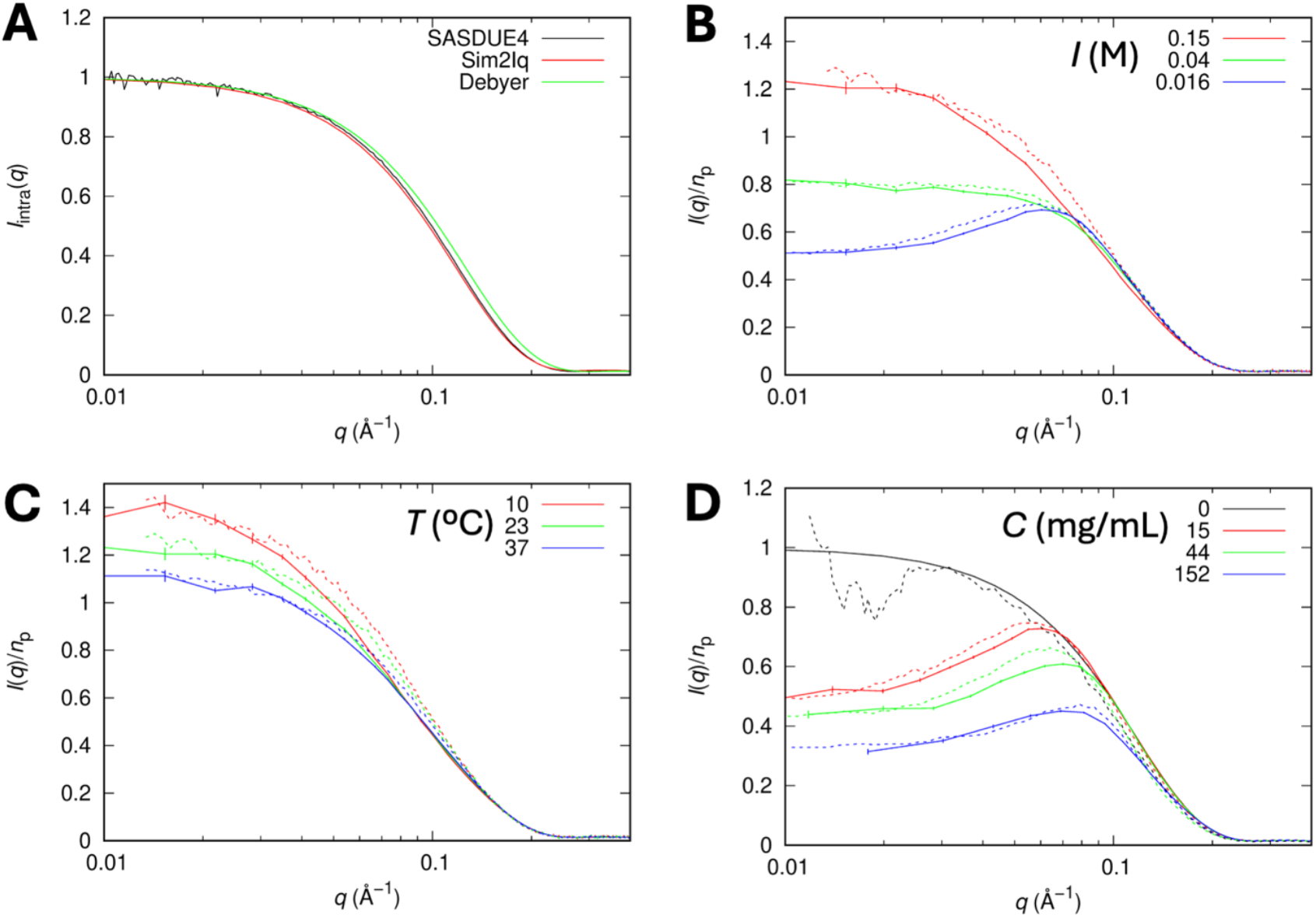
*I*(*q*)/*n*_p_ results for lysozyme. (A) The 0-concentration limit, *I*_intra_(*q*). (B) Results at the indicated ionic strengths, temperatures of 23, 25, and 25 ºC, respectively, and 20 mg/mL concentration. Here and in the remaining figures presenting comparisons between Sim2Iq and experiment, the former curves are solid, with error bars representing the standard error of the mean, and the latter curves are dash. (C) Results at the indicated temperatures and an ionic strength of 0.15 M. Experimental data in (B) and (C) are from Tanouye et al. ^32^. (D) Results at the indicated concentrations (in mg/mL), ionic strengths of 0.015, 0.025, and 0.045 M, respectively, and temperature of 20 ºC. The “0-concentration” curves: *I*_intra_(*q*) from Sim2Iq and data acquired at 1.6 mg/mL. Experimental data are from Shukla et al. ^14^.

In Figure 3B, C, we compare Sim2Iq results for *I*(*q*)/*n*_p_ of 20 mg/mL lysozyme at various ionic strengths and temperature with the experimental data of Tanouye et al. ^32^. The agreement is good for all solvent conditions. Around room temperature, inter-lysozyme interactions are repulsive at low ionic strengths, leading to a value < 1 for *I*(0^+^)/*n*_p_. As the ionic strength is increased to 0.15 M, electrostatic repulsion is weakened and nonpolar attraction takes over, *I*(0^+^)/*n*_p_ increases to above 1 (Figure 3B). For this ionic strength, the effect of attraction is strengthened at a lower temperature (10 ºC) and attenuated at a higher temperature (37 ºC), as reflected by an increase and decrease in *I*(0^+^)/*n*_p_, respectively (Figure 3C).

In Figure 3D, we present the dependence of *I*(*q*)/*n*_p_ on lysozyme concentration at low ionic strengths, where inter-lysozyme interactions are repulsive as just mentioned. With a modest protein concentration-dependent increase in ionic strength to account for ion release, Sim2Iq calculations reproduce well the experimental data collected by Shukla et al. ^14^ at 20 ºC in H_2_O. At increasing protein concentrations, *I*(0^+^)/*n*_p_ decreases progressively, reflecting amplified effects of inter-protein repulsion as the molecules are forced to be closer to each other. Another interesting feature is a peak at intermediate *q*. The peak position, *q*_M_, shifts to higher *q* values as the protein concentration (or number density *n*_p_) increases. This shift can be understood by relating *q*_M_ to the nearest-neighbor distance *d*_nn_ via *q*_M_ ≈ 2π/*d*_nn_ and noting a scaling relation *d*_nn_ ≈ 1.24*n*_p_^−1/3^, leading to

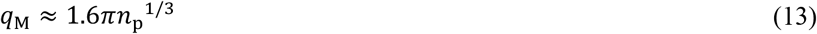

Sim2Iq calculations also reproduce well the experimental data collected by Shukla et al. ^14^ at 10 ºC in D_2_O (Figure S8), when the scaling factor for nonpolar attraction between protein molecules is increased from 0.16 to 0.20 to account for the well-known stabilization effect of D_2_O ^44^. The same qualitative features are observed, namely decrease in *I*(0^+^)/*n*_p_ and rightward shift of the peak position at increasing concentrations.

To investigate more fully the dependences on protein concentration, we carried out Sim2Iq calculations for lysozyme over the concentration range of 10 to 225 mg/mL, under both repulsive (fixed ionic strength at 0.03 M) and attractive (fixed ionic strength at 0.15 M) conditions. Figure S9A presents the *I*(*q*)/*n*_p_ curves for the repulsive case. Consistent with the results shown in Figures 3D and S8, *I*(0^+^)/*n*_p_ decreases with increasing protein concentration. In addition, a prominent peak at intermediate *q* emerges when the concentration reaches 50 mg/mL; the peak position shifts to higher values with further increases in concentration, as predicted by Eq (13) (Figure S9A inset).

*I*(*q*)/*n*_p_ curves for the attractive case are displayed in Figure S9B. As expected, *I*(0^+^)/*n*_p_ is above 1 at 10 mg/mL and increases at 20 mg/mL. However, it becomes obvious that the *I*(*q*)/*n*_p_ curve at 30 mg/mL is no longer a monotonically decreasing function of *q*, with a peak at *q* = 0.018 Å^-1^. The peak height reaches a maximum at 40 mg/mL but then gradually decreases at higher concentrations. On the other hand, the peak positions shift monotonically to higher *q* values with increasing concentrations. The appearance of a peak and its shift to higher *q* values is related to the decrease of *I*(0^+^)/*n*_p_ with increasing concentration (Figure S9B); the decrease of *I*(0^+^)/*n*_p_ in turn is attributed to a decrease in the peak value of *g*(*R*) [see Eqs (2) and (3)]. In Figure S9C, D, we show *g*(*R*) curves, and direct the reader’s attention to the trend of the curves beyond the hard core as the protein concentration increases. For the repulsive case, the *g*(*R*) curves move toward a value of 1 from below, reflecting the tendency for protein molecules to take up all free space at high concentrations. For the attractive case, the *g*(*R*) curves initially move up but then move down toward a value of 1. At increasing concentrations, more and more nearest neighbors localize to the surface of a given protein molecule. However, there is a limit to the number of nearest neighbors that can be closely packed; therefore the increase in the local density cannot keep pace with the increase in overall concentration, leading to a decrease in the peak value of *g*(*R*). Such a decrease has been seen in short-ranged square-well fluids ^45^.

In both the repulsive and attractive cases, the *I*(*q*)/*n*_p_ peak positions shift to higher *q* values at increasing concentrations. However, the peaks in the attractive case occur at much lower *q* values, roughly 50%, of their counterparts in the repulsive case. Whereas the peaks in the repulsive case are associated with dimers formed with nearest neighbors as explained above, the lower-*q* peaks in the attractive case perhaps correspond to larger clusters, which would be separated from each other at a longer distance than *d*_nn_ estimated from the overall concentration. To support this idea, in Figure S9E, F we display the cluster size distributions. The attractive case populates much larger clusters than the repulsive case, e.g., with clusters over 400 molecules compared to 40 molecules at 175 mg/mL. At 225 mg/mL, a very large cluster of ∼700 molecules (out of 800 total) was always formed with voids filled by small clusters and monomers, a sign for imminent liquid-liquid phase separation.

### *S*_eff_(*q*) results for lysozyme

In Figure 4A, we compare Sim2Iq results for *S*_eff_(*q*) of 20 mg/mL lysozyme at various ionic strengths and temperature with the experimental data of Tanouye et al. ^32^. As Sim2Iq predicts *I*(*q*)/*n*_p_ well (Figure 3B, C), the corresponding *S*_eff_(*q*) also reproduces well the experimental counterpart.

**Figure 4.**
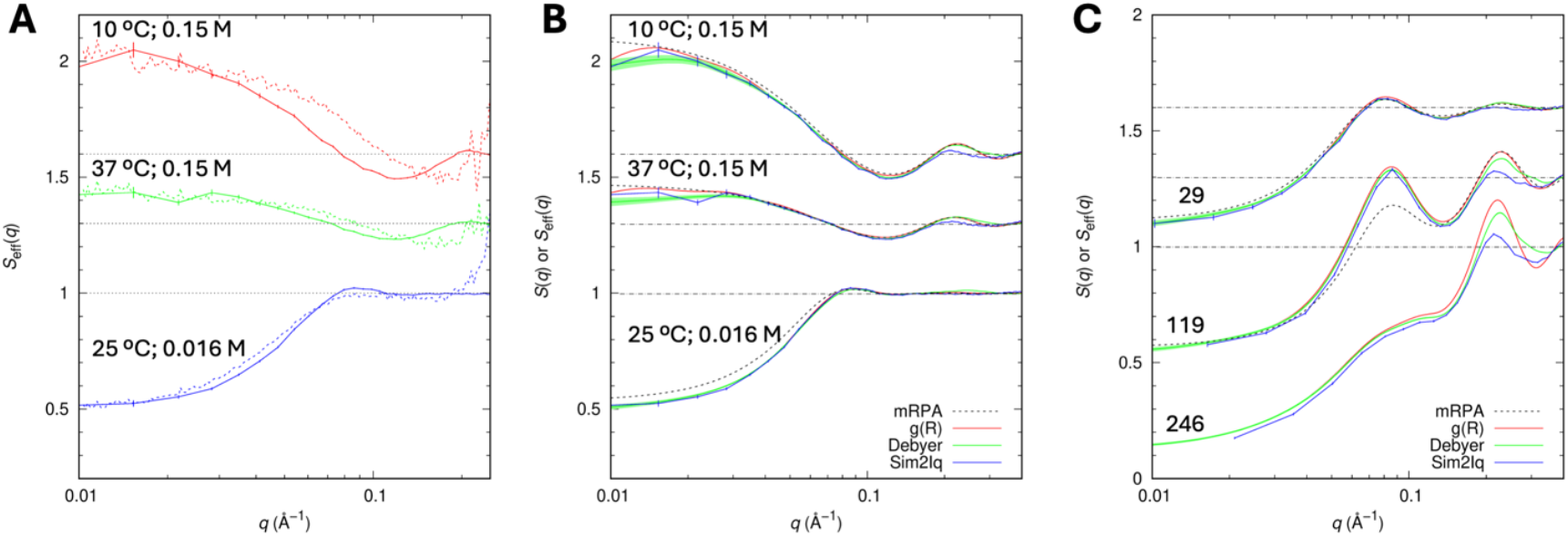
*S*_eff_(*q*) results for lysozyme. (A) Comparison of Sim2Iq and experimental results at the indicated temperatures and ionic strengths; the protein concentration is 20 mg/mL. Experimental data are from Tanouye et al. ^32^. (B) Comparison of the Sim2Iq results in (A) with other *S*_eff_(*q*) or *S*(*q*) calculations. SEM is shown as a band for Debyer. (C) Similar comparison but for the indicated concentration (in mg/mL), ionic strengths of 0.018, 0.024, and 0.055 M, respectively, and temperature of 10 ºC in D_2_O. mRPA breakdowns at 246 mg/mL and is not shown.

We also used the Sim2Iq results to check other calculations, including *S*_eff_(*q*) by Debyer and *S*(*q*) from *g*(*R*) and by mRPA. For lysozyme at 20 mg/mL, all the calculations agree well with each other (Figure 4B). However, *g*(*R*), mRPA, and Debyer all tend to exaggerate the oscillations of *S*_eff_(*q*) at *q* > 0.2 Å^-1^ for attractive conditions (0.15 M ionic strength; 10 or 37 ºC). Even under repulsive conditions (0.018 to 0.055 M ionic strength; 10 ºC), the exaggerated oscillations by *g*(*R*) become very prominent at higher concentrations (Figure 4C). mRPA predicts *g*(*R*) results reasonably well up to ∼100 mg/mL but breaks down at higher concentrations. Exaggerated oscillations are a common problem for all *g*(*R*)-based calculations, which in essence assume spherical models of interprotein interactions, and are also exhibited in *I*(*q*) fits by Shukla et al. using spherical models ^14^. The oscillations in *S*_eff_(*q*) by Debyer are somewhat tempered, due to the atomistic representation of protein molecules. The amplitudes of Sim2Iq *S*_eff_(*q*) oscillations are very small; apparently the inclusion of a hydration layer further blunts the oscillations, as the vacuum-only Sim2Iq results match the Debyer results (Figure S10).

### *I*(*q*) results for BSA

In Figure 5A, we compare Sim2Iq results for *I*(*q*)/*n*_p_ of bovine serum albumin (BSA) at 80-300 mg/mL and low ionic strengths with the experimental data of Zhang et al. ^13^. Similar to results for lysozyme (Figures 3D and S8), a peak appears and shifts to higher *q* at increasing BSA concentration. The peak positions of the calculated *I*(*q*) curves are close to the observed *q* values, but the peak heights are underestimated. Many studies have presented evidence of BSA dimerization ^46^. We thus performed simulations of BSA monomer-dimer mixtures, using a crystal dimer for the structure of the dimer in the simulations. The experimental peak heights are now well reproduced with 10% dimer at 80-200 mg/mL and 40% dimer at 300 mg/mL (Figure 5B). These dimer populations are roughly those predicted by a monomer-dimer equilibrium with a dissociation constant of 550 mg/mL, which is comparable to a value determined by sedimentation equilibrium experiments ^47^.

**Figure 5.**
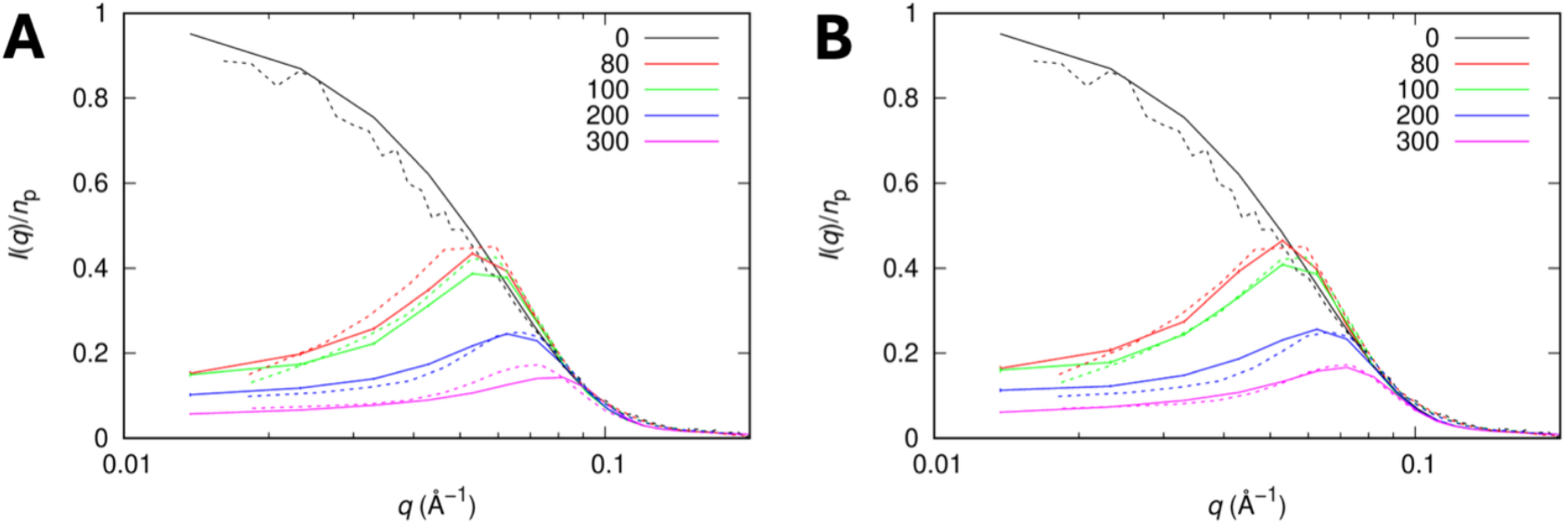
*I*(*q*)/*n*_p_ results for BSA. (A) Comparison of Sim2Iq and experimental results at the indicated concentrations (in mg/mL), ionic strengths of 0.016, 0.02, 0.04, and 0.06 M, respectively, and temperature of 20 ºC. The “0-concentration” curves: *I*_intra_(*q*) from Sim2Iq and data acquired at 10 mg/mL. Experimental data are from Zhang et al. ^13^. (B) Similar comparison but Sim2Iq calculated from simulations with 10% of the molecules at 80 – 200 mg/mL and 40% of the molecules at 300 mg/mL modeled as preformed dimers.

## Discussion

We have presented a fast method for performing simulations of dense protein solutions with an atomistic representation for proteins. From these simulations we directly calculate the small-angle scattering profile *I*(*q*), forgoing the many approximations used up to now. Application of this MAEPPIsim + Sim2Iq approach to lysozyme and BSA demonstrates accurate reproduction of experimental *I*(*q*) data.

The fast calculation of *I*(*q*) enables thorough exploration into the dependences on solvent conditions and protein concentrations, leading to more sound physical interpretations (Figure 6). At low concentrations, *I*(*q*)/*n*_p_ can be regarded as arising from the scattering by a single protein molecule [i.e., *I*_intra_(*q*)], which can be calculated from a crystal structure of the protein. Within the medium concentration range, attraction between protein molecules elevates *I*(0^+^)/*n*_p_ to above 1 whereas repulsion depresses it to below 1; a small increase in concentration accentuates these effects. The behaviors of *I*(*q*)/*n*_p_ at high concentrations are complex and still understudied. Under repulsive conditions, *I*(*q*)/*n*_p_ exhibits a peak at an intermediate *q, q*_M_, that corresponds to the nearest-neighbor distance [see Eq (13)]. Preformed dimers raise this peak whereas preformed higher oligomers elevate the initial rise of the *I*(*q*)/*n*_p_ curve. Under weakly attractive conditions, *I*(*q*)/*n*_p_ might continue the rising trend seen at medium concentrations. However, for stronger attraction that extends only a short range, *I*(*q*)/*n*_p_ again exhibits a peak, but at *q* < *q*_M_ and related to higher oligomers. The latter prediction is yet to be tested.

**Figure 6.**
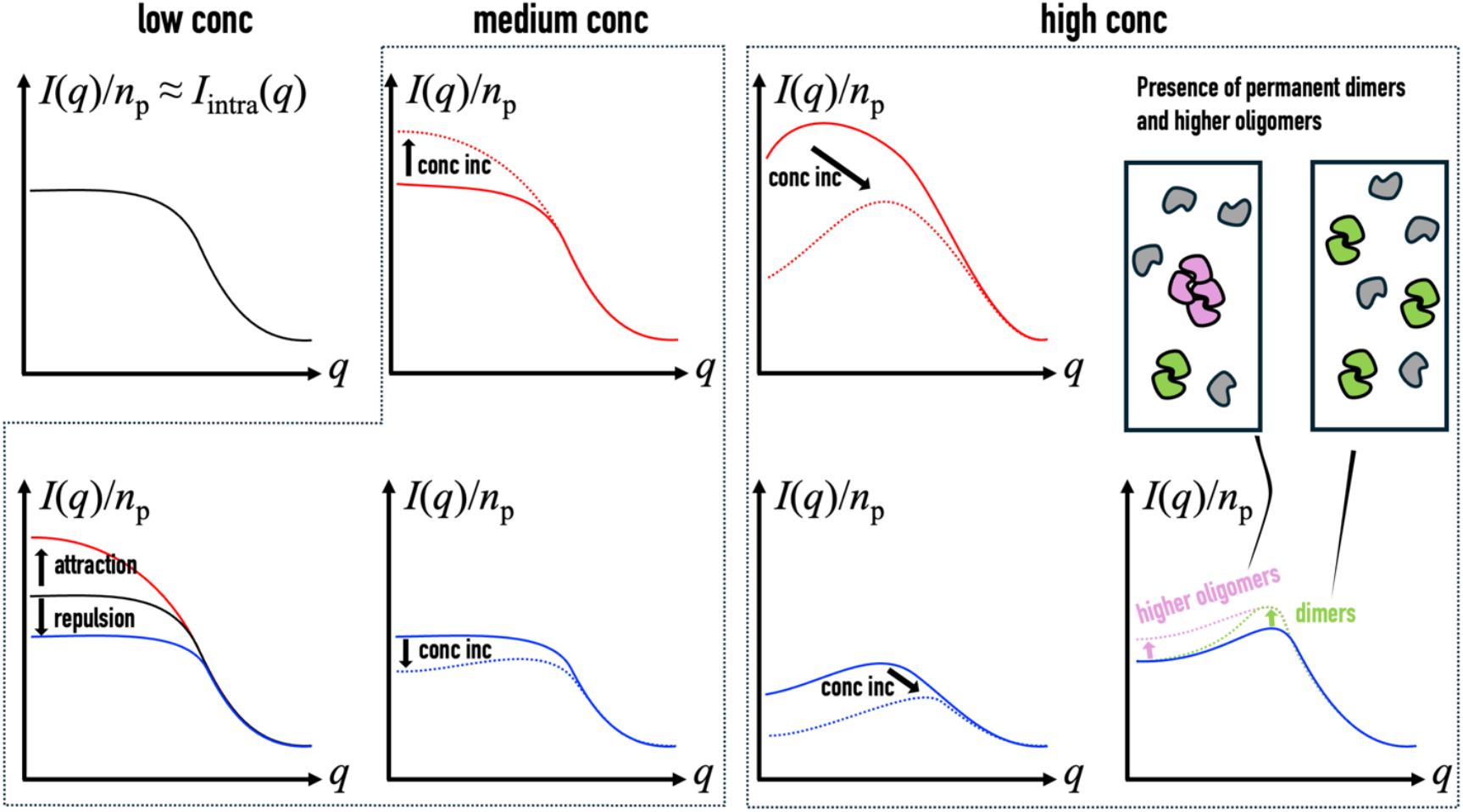
Physical interpretations of *I*(*q*) curves.

The MAEPPIsim + Sim2Iq approach can be further developed on several fronts. (1) Reparameterization of the interaction energy function. Our previous parameterization was based on second virial coefficient data ^38^. The fast speed of *I*(*q*) calculations now makes it practical to use *I*(*q*) data for parameterization. (2) Application to protein mixtures. MAEPPIsim can be readily extended to mixtures, by precomputing interaction energy tables for all unique protein pairs. For binary and ternary mixtures, the number of precomputed tables increases from 1 to 3 and 6, respectively. Sim2Iq is unaffected, as all atoms are mapped to a grid. (3) Accounting for molecular flexibility. Further improvement in accuracy may require a flexible instead of a rigid representation of protein molecules in MAEPPIsim, as that enables them to form more attractive interactions ^40^. These developments promise to modernize the prediction and interpretation of small-angle scattering profiles of dense protein solutions.

### Online Methods

#### Protein pair interaction energies: precomputation and retrieval

We precomputed pair interaction energies at various center-to-center displacements and relative orientations, and stored the data in a “table” (Figure 1B). The table has a 4-dimensional structure, with the first three dimensions for the Cartesian components of the displacement vector **R** (indices running from 0 to blen − 1 per dimension) and the fourth dimension an index for different rotation matrices (index running from 0 to nrot − 1). The discretization of the **R** space used a 0.6-Å spacing in each dimension ^37,38^. To uniformly sample the rotational space, we took the “cardinal” rotation matrices (nrot = 72, 576, or 4608) from Mitchell ^39^.

The pair interaction energy function was the same as described in the original FMAPB2 paper ^38^, except for a modification to the cutoff distance. Instead of a fixed value at 36 Å, the cutoff distance is now the larger of 36 Å and three times the Debye screening length. The scaling factors for nonpolar attraction and electrostatic interactions were changed to account for salt effects and solvent isotope effects (see Table S1).

In a many-protein system, we represented the center position of molecule *i* by a 3-dimensional vector **R**_)_ and its orientation by three Euler angles (*θ*_*i*_, ϕ_*i*_, *ψ*_2_). For a pair of molecules *ij*, the retrieval of their interaction energy required the displacement vector in the body-fixed frame of molecule *i* and the index for the relative rotation matrix. The former was found as

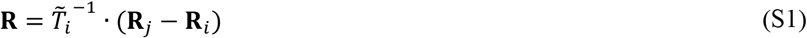

We only stored the nrot cardinal rotation matrices. To map Euler angles to an index for the cardinal rotation matrices, we discretized cos *θ*, ϕ, and *ψ* into bins, with a default value of 120 for the number of bins (nbin). For each of the nbin^3^ sets of Euler angles, we found the nearest cardinal rotation matrix. After mapping the Euler angles of both molecules *i* and *j* to rotation indices, we calculated their relative rotation [see Eq (13)], for which we finally found the nearest cardinal rotation matrix. Both the nbin^3^ → nrot mapping and the nrot^2^ → nrot mapping were precomputed.

### MAEPPI simulations

We implemented MAEPPIsim as Monte Carlo simulations in the canonical ensemble, where the molecule number (*N*), volume (*V*), and temperature (*T*) are held constant. The many-protein system was initialized by sequentially adding molecules into the simulation box. Each new molecule took a random position and orientation, and was placed repeatedly until free of clash with any existing molecules. A Monte Carlo cycle consisted of a sweep of *N* steps. At each step, a molecule was randomly selected for a trial move in translational and rotational space. A new position in translational space was randomly selected inside a cubic box that was centered at the old position and had a side length adjusted according to the number density. A new position in rotational space defined by cos *θ*, ϕ, and *ψ* was similarly selected from a cubic box with a fixed side length. Periodic boundary conditions were enforced in both translational and rotational space: for the former, Cartesian coordinates were wrapped into the simulation box; for the latter, cos *θ* was wrapped into [−1,1] whereas ϕ, and *ψ* were wrapped into [−π, π]. The move was accepted or rejected based on the Metropolis criterion: a move that lowered the energy was always accepted, while a move that increased the energy by Δ𝕌 was accepted with a probability exp (−Δ𝕌 /*k*_B_*T*), where *k*_B_ is the Boltzmann constant. For each trial move, Δ𝕌 was calculated using only the pairs formed between the selected molecule and the rest of the molecules.

Each Monte Carlo simulation was run for 20,000 cycles. Calculations were averaged over 64 replicate simulations. Error bars represented the standard error of the mean.

### Sim2Iq calculations

Sim2Iq calculates the small-angle X-ray scattering intensity profile *I*(*q*) from atomic coordinates of protein molecules in a periodic box. It reimplements the core functionality of DENSS ^26,27^ but with two major changes. First, it handles periodic boundary conditions so that not only *I*_intra_(*q*) but also *I*(*q*) at arbitrary protein concentrations can be calculated. Second, it uses OpenMP for parallelization to achieve dramatic speedup.

Sim2Iq is similar in spirit to FMAP^36,37^ by going to Fourier space and taking advantage of the speed of FFT. In direct space, the excess electron density consists of three terms [see Eq (10)]. Atomic representations of these terms were mapped to a cubic grid, with periodic boundary conditions enforced. After FFT into **q** space, spherical averaging and binning produced *I*(*q*) at discrete *q* values.

## Acknowledgment

We sincerely thank Prof. Rosangela Itri for sharing experimental data and providing clarifications under challenging circumstances. This work was supported by National Institutes of Health Grant GM118091.

## Notes

### Competing Interest Statement

The authors have declared no competing interest.

