## Supplementary Table and Figures for "Fast calculation of small-angle scattering profiles of dense protein solutions modeled at the all-atom level"

*Supplementary Information*

Table S1. Experimental conditions and calculation parameters.

| Figure # | Expt. conditions (Temp; buffer) | Conc (mg/mL) | Ionic strength (M) | Net charge | Precomputing | | MAEPPIsim[a] | |
|---|---|---|---|---|---|---|---|---|
| | | | | | # grids per dim | # of rotations | # of mol. | Box side length (Å) |
| 2 | | 20 | 0.15 | 8 | 310 | 72 | 800 | 984.15 |
| 3B | 25 ºC, 10 mM Tris pH 8 | 20 | 0.016 | 7 | 436 | 72 | 800 | 984.15 |
| | 25 ºC, 10 mM Tris pH 8 + 50 mM NaCl | 20 | 0.04 | 7 | 326 | 72 | 800 | 984.15 |
| | 23 ºC, PBS pH 7.4[b] | 20 | 0.15 | 8 | 310 | 72 | 800 | 984.15 |
| 3C | 10/23/37 ºC, PBS pH 7.4[b] | 20 | 0.15 | 8 | 310 | 72 | 800 | 984.15 |
| 3D | 20 ºC, 20 mM Hepes pH 7.8 | 15 | 0.015 | 7 | 466 | 72 | 800 | 1081.35 |
| | | 44 | 0.025[c] | 7 | 382 | 72 | 800 | 757.35 |
| | | 152 | 0.045[c] | 7 | 332 | 72 | 800 | 498.15 |
| 4A,B | | Same as 3B,C | | | | | | |
| 4C | | 29 | 0.018 | 7 | 420 | 72 | 800 | 870.75 |
| | | 119 | 0.024 | 7 | 386 | 72 | 800 | 543.7 |
| | | 246 | 0.055 | 7 | 320 | 72 | 800 | 425.25 |
| 5 | 20 ºC, no buffer, pH 7 | 80 | 0.016[c] | -15 | 578 | 72 | 200 | 648 |
| | | 100 | 0.02[c] | -15 | 554 | 72 | 248 | 648 |
| | | 200 | 0.04[c] | -15 | 492 | 72 | 496 | 648 |
| | | 300 | 0.06[c] | -15 | 462 | 72 | 744 | 648 |
| S1 | | 100 | 0.15 | 8 | 310 | 72 | 200 | 362.14 |
| | | 100 | 0.15 | 8 | 310 | 72 | 400 | 456.26 |
| | | 100 | 0.15 | 8 | 310 | 72 | 600 | 522.29 |
| | | 100 | 0.15 | 8 | 310 | 72 | 800 | 574.86 |
| | | 50 | 0.15 | 8 | 310 | 72 | 200 | 456.26 |
| | | 100 | 0.15 | 8 | 310 | 72 | 400 | 456.26 |
| | | 150 | 0.15 | 8 | 310 | 72 | 600 | 456.26 |
| | | 200 | 0.15 | 8 | 310 | 72 | 800 | 456.26 |
| S2A | | 225 | 0.065 | 10 | 310 | 72/576/ 4608 | 800 | 437.4 |
| S2B-D | | 225 | 0.065 | 10 | 310 | 72/576/ 4608 | 322 | 324 |
| S3 | | 70 | 0.15 | 8 | 310 | 72 | 100 | 324 |
| S4 | | 20 | 0.016 | 7 | 436 | 72 | 800 | 984.15 |
| S5A | | Same as S2B-D | | | | | | |
| S5B | | 20 | 0.016 | 7 | 436 | 72 | 800 | 984.15 |
| | | 20 | 0.016 | 7 | 436 | 72 | 400 | 781.65 |
| | | 20 | 0.016 | 7 | 436 | 72 | 200 | 619.65 |
| S5C | | 20 | 0.04 | 7 | 326 | 72 | 800 | 984.15 |
| | | 20 | 0.04 | 7 | 326 | 72 | 400 | 781.65 |
| | | 20 | 0.04 | 7 | 326 | 72 | 200 | 619.65 |
| S5D | | 20 | 0.15 | 8 | 310 | 72 | 800 | 984.15 |
| | | 20 | 0.15 | 8 | 310 | 72 | 400 | 781.65 |
| | | 20 | 0.15 | 8 | 310 | 72 | 200 | 619.65 |
| S7 | | Same as 3B | | | | | | |
| S8 | 10 ºC, 20 mM Hepes pH 7.8[d] | 29 | 0.018[c] | 7 | 420 | 72 | 800 | 870.75 |
| | | 119 | 0.024[c] | 7 | 386 | 72 | 800 | 543.7 |
| | | 246 | 0.055[c] | 7 | 320 | 72 | 800 | 425.25 |
| S9A,C,E | | 10 | 0.03 | 8 | 366 | 72 | 400 | 984.15 |

| | | | | | | | | |
|---|---|---|---|---|---|---|---|---|
| | | 20 | 0.03 | 8 | 366 | 72 | 800 | 984.15 |
| | | 30 | 0.03 | 8 | 366 | 72 | 800 | 858.6 |
| | | 40 | 0.03 | 8 | 366 | 72 | 800 | 781.65 |
| | | 50 | 0.03 | 8 | 366 | 72 | 800 | 724.95 |
| | | 100 | 0.03 | 8 | 366 | 72 | 800 | 575.1 |
| | | 150 | 0.03 | 8 | 366 | 72 | 800 | 502.2 |
| | | 175 | 0.03 | 8 | 366 | 72 | 800 | 477.9 |
| | | 225 | 0.03 | 8 | 366 | 72 | 800 | 437.4 |
| S9B,D,F | | 10 | 0.15 | 8 | 310 | 72 | 400 | 984.15 |
| | | 20 | 0.15 | 8 | 310 | 72 | 800 | 984.15 |
| | | 30 | 0.15 | 8 | 310 | 72 | 800 | 858.6 |
| | | 40 | 0.15 | 8 | 310 | 72 | 800 | 781.65 |
| | | 50 | 0.15 | 8 | 310 | 72 | 800 | 724.95 |
| | | 100 | 0.15 | 8 | 310 | 72 | 800 | 575.1 |
| | | 150 | 0.15 | 8 | 310 | 72 | 800 | 502.2 |
| | | 175 | 0.15 | 8 | 310 | 72 | 800 | 477.9 |
| | | 225 | 0.15 | 8 | 310 | 72 | 800 | 437.4 |
| S10 | | Same as 4B,C | | | | | | |

[a]In all cases, MAEPPIsim ran for 20,000 Monte Carlo cycles. All Sim2Iq calculations used the final snapshot; for Figure 2B, Sim2Iq calculations were also carried out for the initial snapshot and the snapshot after 2,000 Monte Carlo cycles. Sim2Iq calculations were done using a 1-Å spacing for mapping excess electron density to a cubic grid; additional calculations using a 0.5-Å spacing for Figure S4A, B. All $g(R)$ and cluster calculations used all the snapshots in the last 2,000 Monte Carlo cycles. Results were averaged over 64 replicate simulations, except for Figure S2B, C, where a single simulation was used.

[b]PBS buffer contains 137 mM NaCl and 2.7 mM KCl. We scaled down electrostatic interactions by a factor of 0.4 to account for anion binding, and scaled up nonplar attraction by increasing its weighting factor from 0.16 to 0.168. These modifications apply to all simulations at an ionic strength of 0.15 M.

[c]To account for ion release from protein surface, ionic strength was increased a small amount, roughly 0.02 M per 100 mg/mL of protein.

[d]This experiment was done in $D_2O$. To account for the stabilization effect of $D_2O$, we scaled up nonplar attraction by increasing its weighting factor from 0.16 to 0.2.

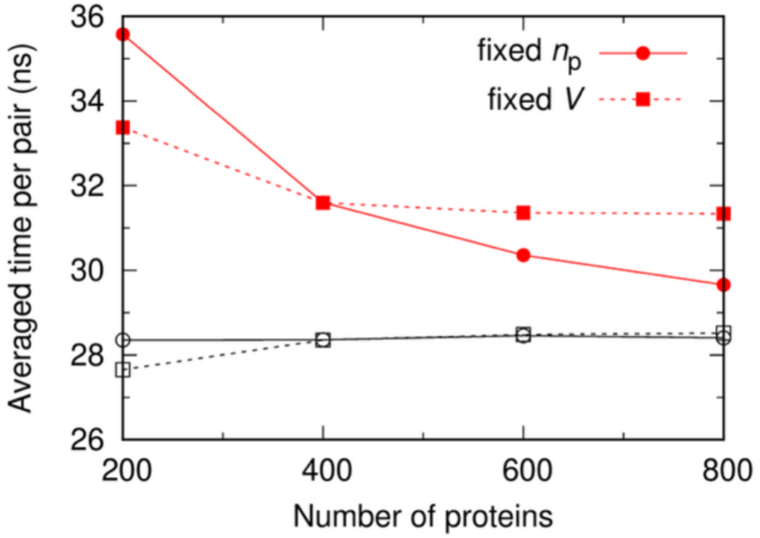

Figure S1. Comparison of CPU times for MAEPPIsim and corresponding simulations of Lennard-Jones particles. For the fixed $V$ series, the simulation boxes have a side length of 456 Å (concentrations at 50, 100, 150 and 200 mg/mL); for the fixed $n_p$ series, the concentration is 100 mg/mL, and box side lengths are 362, 456, 522, and 575 Å. Solvent conditions: 10 ºC and 0.15 M ionic strength.

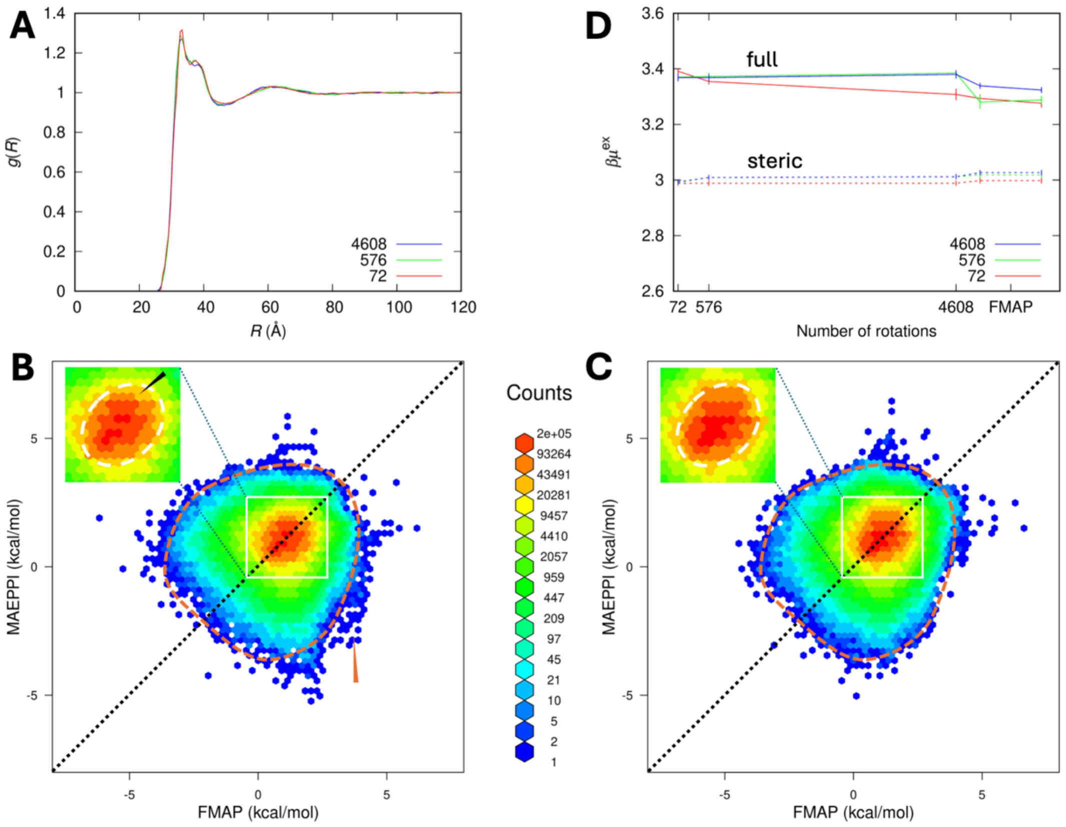

Figure S2. Benchmarking of MAEPPI. (A) $g(R)$ results from MAEPPIsim using different numbers of rotations for precomputing pair interactions. The simulation box had a side length of 437 Å and contained 800 lysozyme molecules (corresponding to 227 mg/mL). Solvent conditions: 25 ºC and 0.065 M ionic strength. (B) Comparison of MAEPPI and FMAP energy, for the insertion of a test protein into a MAEPPIsim final snapshot, at all points on a cubic grid. The simulation box had a side length of 324 Å and contained 322 lysozyme molecules (corresponding to 225 mg/mL); MAEPPI used 72 rotations for precomputing pair interactions. An orange arrowhead indicates extra off-diagonal outliers; a black arrow in the inset indicates a lower count of closely matched energies between MAEPPI and FMAP. (C) Similar to (B), but for 4608 rotations. (D) Excess chemical potential, calculated from insertion energies such as shown in (B) and (C). For MAEPPIsim using a given number of rotations (shown in legend), insertion energies were calculated using tables precomputed with three different numbers of rotations (shown in abscissa). For FMAP, the excess chemical potential was calculated in two ways, treating the protein solution and the test protein as "source charge" and "test charge", respectively, or vice versa.

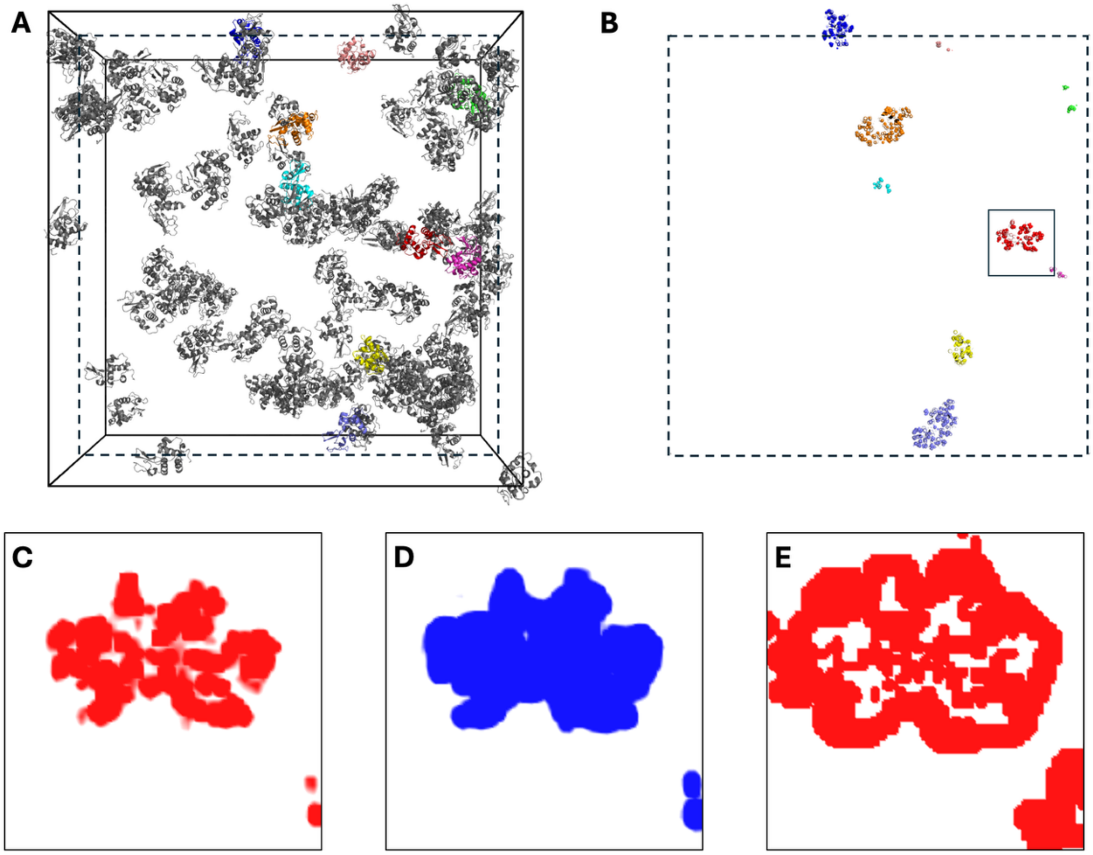

Figure S3. Illustration of mapping the excess electron density to a cubic grid. (A) A simulation box with a side length of 324 Å and containing 100 lysozyme molecules. A slice through the box creates a cross section defined by the dashed square; the intersecting molecules are shown in color. (B) The resulting cross section. A small region of interest (ROI) is indicated by a solid square. (C) The vacuum component of the excess electron density in ROI. (D) The solvent electron density in the region excluded by protein molecules. (E) The hydration shell.

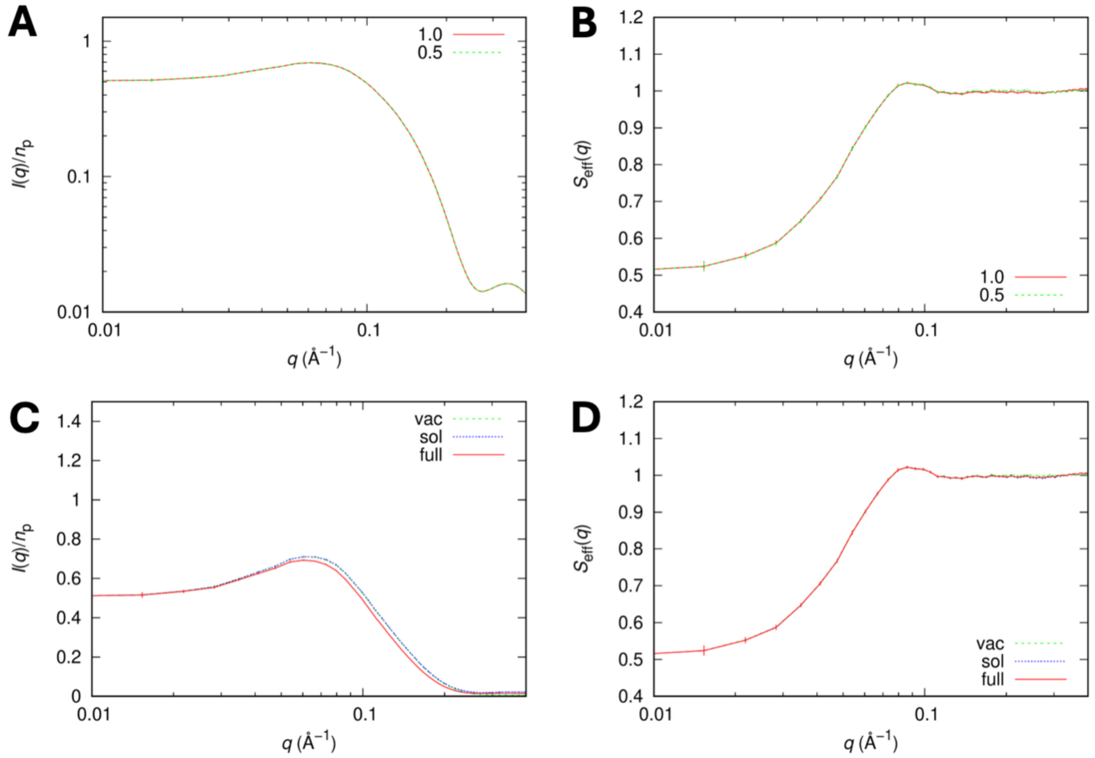

Figure S4. Effects of grid spacing and the different components of the excess electron density on calculated $I(q)$ and $S_{\text{eff}}(q)$. (A) Comparison of $I(q)$ results calculated using 1.0 and 0.5 Å grid spacing in mapping the excess electron density. (B) Corresponding comparison for $S_{\text{eff}}(q)$. (C) $I(q)$ results when only the vacuum component ("vac"), both the vacuum and solvent-exclusion components ("sol"), and the full excess electron density ("full") are included. (D) Corresponding comparison for $S_{\text{eff}}(q)$. The MAEPPIsim box had a side length of 984 Å and contained 800 lysozyme molecules. Solvent conditions: 25 ºC and 0.016 M ionic strength.

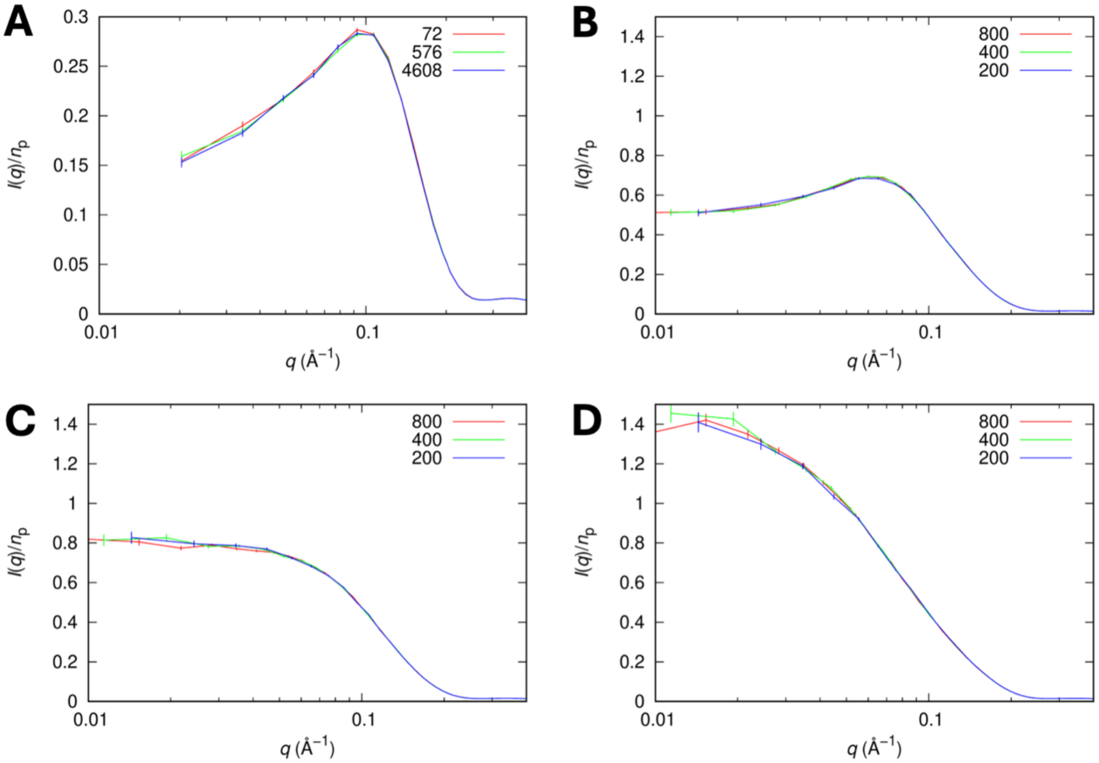

Figure S5. Further benchmarking of $I(q)$. (A) Comparison of results calculated from MAEPPIsim using different numbers of rotations. The simulation box had a side length of 437 Å and contained 800 lysozyme molecules (corresponding to 227 mg/mL). Solvent conditions: 25 ºC and 0.065 M ionic strength. (B-D) Comparison of results calculated from MAEPPIsim using different numbers of protein molecules for a fixed concentration of 20 mg/mL. The simulation boxes had side lengths of 984, 782, and 620 Å, respectively. Solvent conditions: (B) 25 ºC and 0.016 M ionic strength; (C) 25 ºC and 0.04 M ionic strength; and (D) 10 ºC and 0.15 M ionic strength.

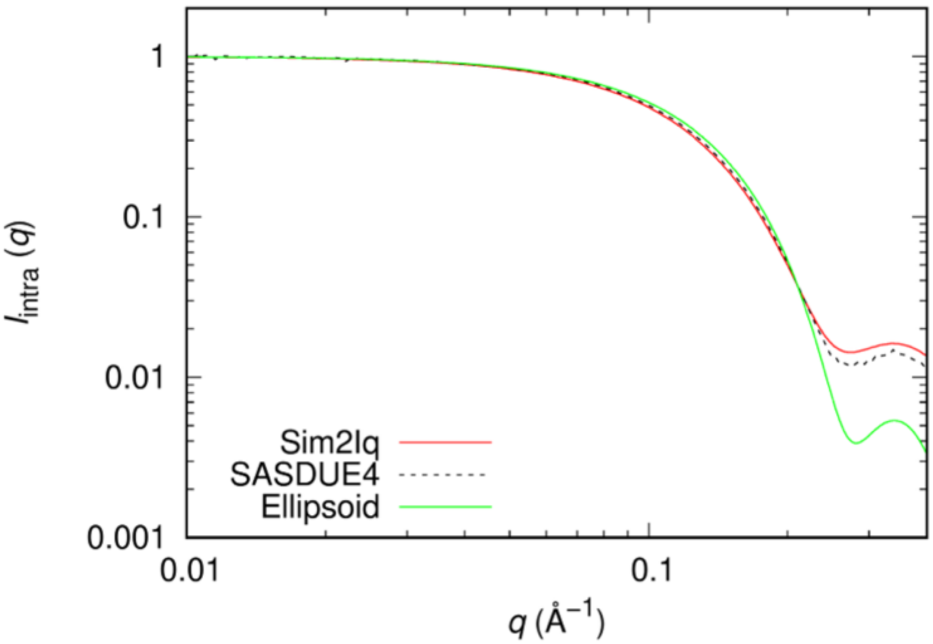

Figure S6. Comparison of $I_{\mathrm{intra}}(q)$ from Sim2Iq, an ellipsoid model, and experiment. Sim2Iq used PDB entry 1AKI as input; the ellipsoid model has semi-axes of 15, 15, and 22.5 Å; the experimental data is from SASBDB entry SASDU4.

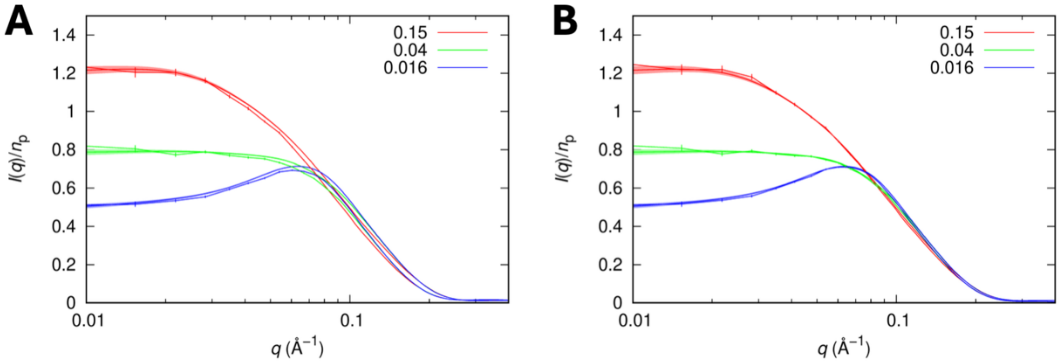

Figure S7. Comparison of Sim2Iq and Debyer results for lysozyme $I(q)/n_{\mathrm{p}}$. (A) Sim2Iq vs, Debyer, with SEMs shown as error bars and a band, respectively. (B) Similar comparison, but Sim2Iq is vacuum only. The simulation box had a side length of 984 Å and contained 800 lysozyme molecules (corresponding to 20 mg/mL). Solvent conditions: ionic strengths at indicated values in M and temperatures at 23, 25, and 25 ºC, respectively.

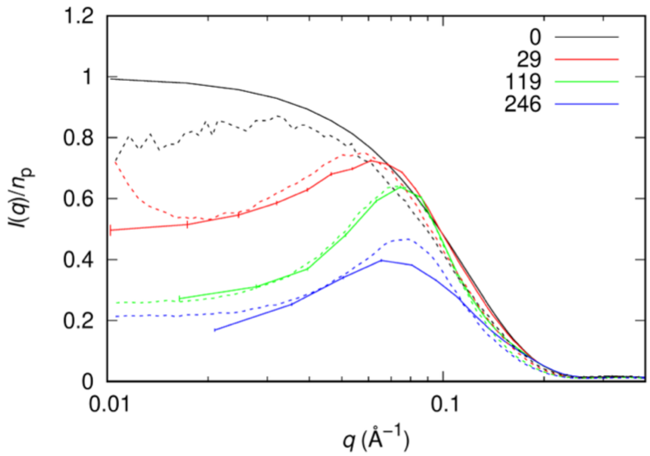

Figure S8. Comparison of Sim2Iq and experimental results for lysozyme $I(q)/n_\mathrm{p}$ in $D_2O$. The protein concentrations are indicated in the legend (in mg/mL); ionic strengths are 0.018, 0.024, and 0.055 M, respectively; temperature = 10 ºC. The “0-concentration” curves: $I_\mathrm{intra}(q)$ from Sim2Iq and data acquired at 4.2 mg/mL. Experimental data are from Shukla et al.

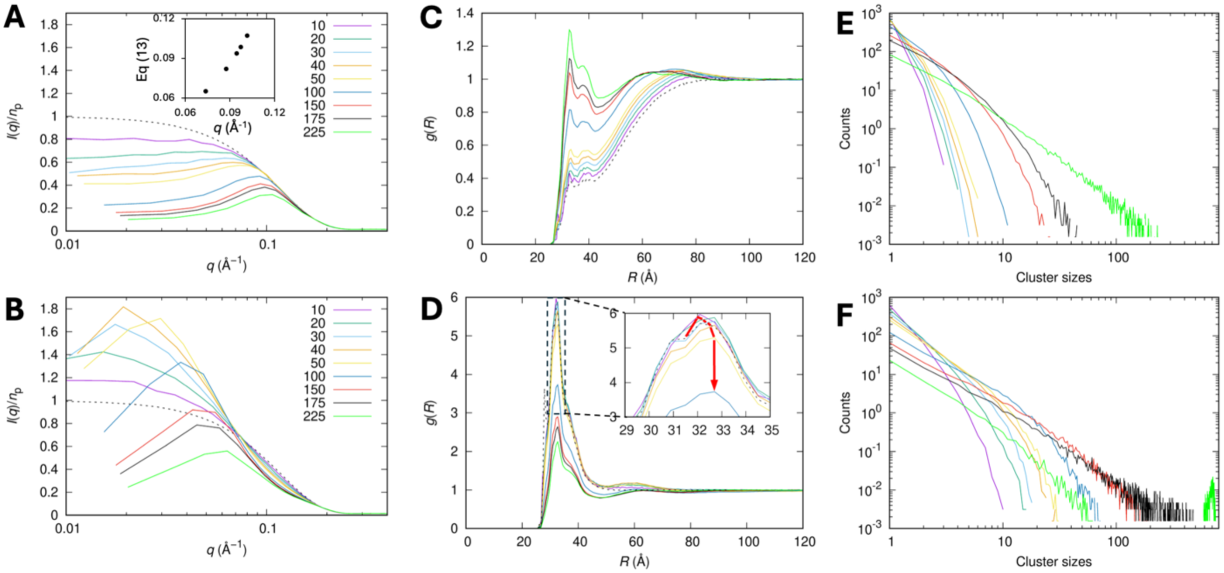

Figure S9. $I(q)/n_\mathrm{p}$, $g(R)$, and cluster histograms for lysozyme. (A, C, E) $I(q)/n_\mathrm{p}$, $g(R)$, and cluster histogram for a repulsive case at the indicated concentrations (in mg/mL). Inset in (A): comparison of the observed $I(q)$ peak positions and those predicted by Eq (13). (B, D, F) Counterparts for an attractive case. Inset in (D): zoomed view of the $g(R)$ peaks; the red, arrowed curve indicates the change in peak height at increasing concentrations. Clusters were calculated based on a center-center cutoff distance of 40 Å. The simulation boxes contained 800 lysozyme molecules at all concentrations except for 10 mg/mL, where the number of molecules was reduced to 400. Solvent conditions: (A, C, E) 25 ºC and 0.03 M ionic strength; (B, D, F) 10 ºC and 0.15 M ionic strength.

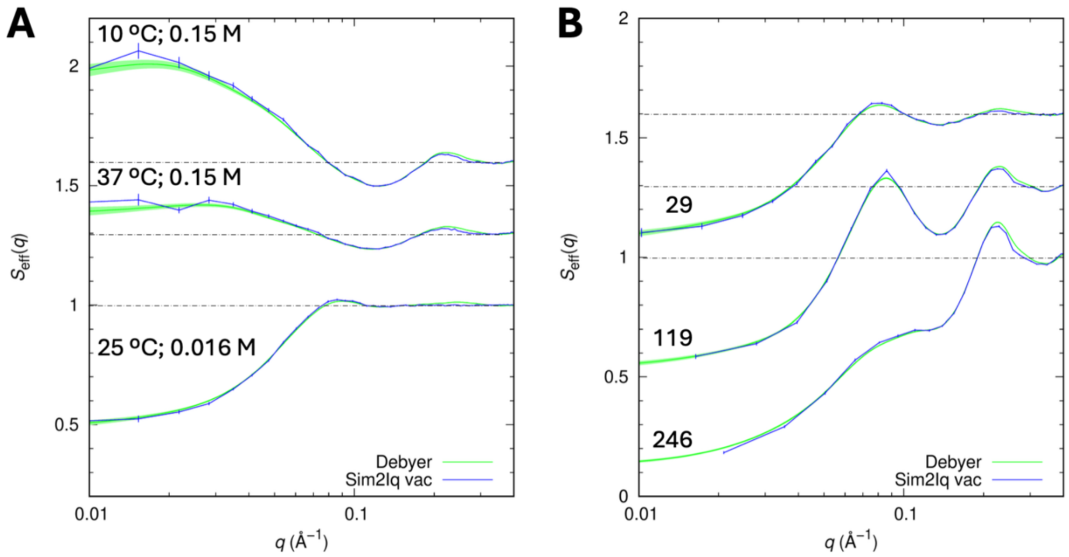

Figure S10. Comparison of lysozyme $S_{\text{eff}}(q)$ results from vacuum-only Sim2Iq and from Debyer. (A) Results at the indicated temperatures and ionic strengths; the protein concentration is 20 mg/mL. (B) Results at the indicated concentration (in mg/mL), ionic strengths of 0.018, 0.024, and 0.055 M, respectively, and temperature of 10 ºC.
